# Single cell genomics and fluorescence microscopy suggest a permanent plastid in a marine centrohelid

**DOI:** 10.64898/2026.06.22.732937

**Authors:** Anne Walraven, Vasily Zlatogursky, Patrick J. Keeling, Rachel A. Foster, Fabien Burki

**Affiliations:** Department of Organismal Biology, Program in Systematic Biology, Uppsala University, Uppsala, Sweden; Botany Department, University of British Columbia, 6270 University Blvd. Vancouver, BC, Canada; Department of Ecology, Environment, and Plant Sciences, Stockholm University, Stockholm Sweden

**Keywords:** plastid evolution, endosymbiosis, horizontal gene transfer, centrohelids

## Abstract

Photosynthetic organelles (plastids), which originated via primary endosymbiosis between an Archaeplastida ancestor and cyanobacteria, have shaped the Earth’s oxygen-rich atmosphere and spread across the eukaryotic tree of life through multiple secondary and higher order endosymbioses. Yet, the mechanisms driving the transition from endosymbiont to organelle remain poorly understood, highlighting the need for novel systems to elucidate the stages of endosymbiont integration and test the generality of plastid evolution models. Here, we employ single-cell genomics and catalyzed reporter deposition fluorescence in-situ hybridization (CARD-FISH) to investigate the dictyochophyte plastids of the poorly known marine centrohelid *Meringosphaera* across diverse geographic locations. Our analyses detected novel microdiversity of cells both containing and lacking plastids. Using phylogenomics we show that host and plastid data are perfectly congruent across a large plastid-containing clade (named MER-2), strongly indicating co-evolution. Extensive environmental screening using double CARD-FISH simultaneously targeting host and plastid further confirms that MER-2 cells almost always harbor plastids, supporting the hypothesis of permanent plastid integration and vertical transmission. Additionally, we find that both MER-2 hosts and other *Meringosphaera* lineages encode multiple plastid-associated genes from diverse phylogenetic origins, with many of their products predicted to be plastid-targeted. Collectively, our findings represent the first report of algae in centrohelids, a large eukaryotic group of otherwise heterotrophic predators. The discovery of new plastids is rare and underscores the importance of exploring uncultured algae to provide new insights into plastid origin and evolution.

## Introduction

The acquisition of organelles through endosymbiosis with bacteria (e.g. mitochondria, plastids, nitroplasts) has been a fundamental process for the evolution and diversification of eukaryotes (1–7). The endosymbiotic origin of plastids remains particularly enigmatic, owing not only to their ancientness but also to their convoluted evolution. Plastids first evolved by primary endosymbiosis with cyanobacteria in an ancestor of Archaeplastida, a supergroup of eukaryotes including the major photosynthetic lineages green algae (and later land plants) and red algae, as well as the smaller group glaucophytes, all containing primary plastids bound by two membranes. Most algal groups, however, do not directly derive from this initial plastid endosymbiosis but instead acquired their plastid by multiple rounds of eukaryote-to-eukaryote endosymbiosis. Both green algae and red algae donated their plastids in these higher order endosymbioses, resulting in more complex plastids with additional membranes and, in some cases, still possessing a highly reduced version of the endosymbiont nucleus (called a nucleomorph).

The separate origin of complex green plastids in euglenids, chlorarachniophytes, and some dinoflagellates is relatively clear, but the origin and evolution of red algal-derived complex plastids is still debated (8–14). Red algal plastids are currently known in cryptophytes, leptophytes, haptophytes, ochrophytes (photosynthetic stramenopiles), and myzozoans (photosynthetic alveolates). Although these plastids have been suggested to be of common secondary origin, host phylogenies do not support a single origin and they have accordingly been proposed to be the result of serial endosymbioses, or even multiple secondary endosymbioses, instead (8, 9, 13–15). Notwithstanding this debate, it is clear that some lineages in dinoflagellates (alveolates) acquired (or re-acquired) plastids from other algae with complex red plastids. The two best known examples are the Kareniaceae with plastids of haptophyte origin (16–19), and the dinotoms with plastids of diatom origin (20–22). While this accounts for most plastid diversity in eukaryotes, there is one significant exception in the cercozoan amoeba *Paulinella*, which has been shown to possess a primary plastid derived from a completely independent endosymbiotic event with a different lineage of cyanobacteria (23, 24). Moreover, in addition to these permanent plastids, there are also many eukaryotes that have gained access to photosynthesis through transient symbiotic associations with algae or cyanobacteria (i.e. photosymbiosis) or through the theft of plastids and sometimes other organelles from algal prey (i.e. kleptoplastidy) (25–27). These associations can be inherited vertically, but not over long evolutionary time scales, and the photosynthetic components need to be refreshed regularly by taking up new algae (28–30).

There are two main hypotheses to explain how endosymbionts transform into permanently integrated and vertically inherited plastids. Traditionally, a linear process has been envisioned where a heterotrophic host engulfed and retained a photosynthetic prey, and over time the endosymbiotic “prey” reduced into a *bona fide* organelle (1). In this model, the genes move from the endosymbiont to the host, and their protein products are retargeted to the plastid where they function, so plastid-targeted proteins derive almost exclusively from the same taxonomic source as the endosymbiont. The alternative model sees the transition to organelles as non-linear. The heterotrophic host feeds on various prey, and over time begins to transiently retain some prey cells for longer periods before digesting them. There are different alternatives of this model, generally referred to as “cyclic” or “shopping bag” models, because the host “shops around” for plastids before eventually fixing one endosymbiont (31, 32). Here, the host evolves a protein-targeting system and gradually acquires genes from various prey and endosymbionts over time, mostly not from the endosymbiont that is ultimately fixed. These two kinds of models predict different phylogenetic origins of plastid-targeted proteins: in the linear model they are predicted to mostly be related to the organelle lineage, and in the cyclic model many genes will be derived from a variety of other lineages. Both kinds of models are consistent with kleptoplastidy as an intermediate stage in complex plastid establishment, but they have different predictions about the order of events and how the host might interact with kleptoplasts (33).

Only a few plastid symbiosis systems have been studied at the level of detail required to distinguish between models, and so far all support a cyclic shopping bag model. The Ross Sea Dinoflagellates (RSD), for example, which steal plastids for long-term survival from haptophyte prey, obtained genes for kleptoplast-targeted proteins by horizontal gene transfers (HGTs) from sources other than the kleptoplast lineage (18). Some of these HGTs are also found in the related species with fully integrated haptophyte plastids, which provided one of the first direct evidence of genetic integration preceding organelle fixation (17, 18). Similar findings were reported in the euglenid *Rapaza viridis*, which is sister to photosynthetic euglenids but itself acquired photosynthesis via kleptoplastidy (29). *R. viridis* specifically steals green algal plastids from a single strain of *Tetraselmis*, yet it harbors genes from a variety of other algae (not just its kleptoplast source) and shares many of these genes with photosynthetic euglenids, pointing to a history of HGTs from diverse algal prey. *Paulinella* and other non-plastid endosymbionts with genetic integration also exhibit this pattern (34–36). However, it remains unclear whether other overlooked algal groups, unrelated to known photosynthetic lineages, have recently acquired plastids–and if so, whether they follow a similar model.

One poorly studied group with plastid endosymbionts is *Meringosphaera* (37). This genus is ubiquitous in marine planktonic samples, especially in the ocean surface, but is generally rare.

The cells can, however, reach high relative abundance under unknown conditions, as attested by two dominating undescribed ASVs that are now attributed to *Meringosphaera* (Centrohelid-sp1 and Centrohelid-sp2 in Obiol et al. (38)). Few historical observations have consistently–although not always (39)–noted that *Meringosphaera* harbours internal autofluorescent green bodies resembling plastids (40–46). This is unexpected based on the phylogenetic placement of *Meringosphaera*, because it belongs to the centrohelids, a relatively small group of heliozoan predatory heterotrophic protists (47). Previously, it was reported that *Meringosphaera* contains two separate groups (group-1 and -2) both with plastids of dictyochophyte (ochrophyte) origin (37). The genomes of these plastids are slightly reduced compared to available dictyochophyte plastid genomes, but appear functional, notably retaining the core repertoire of photosynthetic and carbon fixation genes. Although the exact nature of these plastids remained ambiguous, it was suggested that group-2 harbours kleptoplasts based on a combination of observations: i) No endosymbiont nuclear data was detected in the analysis of 15 single-amplified genomes (SAGs), indicating that the autofluorescing bodies are not full endosymbiotic cells; ii) The plastid signal was below detection during several months in a yearly qPCR survey, while the *Meringosphaera* host remained present; iii) Catalyzed Reporter Deposition - Fluorescence *in situ* Hybridization (CARD-FISH) data of spring samples collected when no qPCR plastid signal was detected showed clear chlorophyll autofluorescence in positively hybridised cells, which was interpreted as evidence for plastid switching (although this second “spring plastid” remained unidentified). It was also found that the host genome of *Meringosphaera* contains some genes that typically function in plastids, some with plastid targeting signal, and of various phylogenetic origins (37). These results provided evidence for gene transfers and protein re-targeting reminiscent of kleptoplastidic systems such as RSD and *R. viridis* (17, 18, 29), and supported the shopping bag model of plastid origin where genetic integration occurs relatively early in endosymbiotic relationships (31).

In this study, we provide extensive SAG and double CARD-FISH data from geographically distinct regions to characterise in detail the diversity and plastid occurrence in *Meringosphaera*. We recovered new diversity for both host and plastid, which now contain six plastid-containing groups resolving as fully mirroring by phylogenomic analyses. A broad survey of environmental samples revealed that nearly all *Meringosphaera* cells belonging to group-2 have clear visual evidence of plastids, but we also identified a plastid-lacking group that branches as sister to group-2. In addition, we searched for host-encoded plastid-associated genes across this new diversity, revealing extensive genetic integration in plastid-containing groups. However, most of these genes appear to have been acquired from multiple sources before *Meringosphaera* presumably obtained its plastids. Collectively, our results support the new interpretation that *Meringosphaera* group-2 harbors a permanent plastid, which makes it the first algal group in centrohelids to have transitioned to photoautotrophy and represents one of very few discoveries of new algae outside of the main established photosynthetic groups.

## Results and Discussion

### 50 new single cell assembled genomes from varied geographic locations reveal novel key diversity of *Meringosphaera*

To expand the known diversity of *Meringosphaera* and its plastids, we manually isolated single cells from three geographically distinct locations and multiple sampling sites (Dataset S1). While sampling in Canada and Curaçao was done at single time points, samples from Sweden were obtained over several months in 2022-2023, including during the months of January to June to target the unknown “spring plastid” previously identified (Dataset S1) (37). After multiple displacement amplification (MDA) and confirmation of *Meringosphaera* presence by 18S rDNA sequencing, we selected a total of 50 single-cells for high-throughput sequencing (Dataset S1).

After assembly into SAGs, we recovered at least partial 18S rDNA (1225-1720 bp) and/or 28S rDNA (308-3016 bp) sequences from 40 single-cells and used these to infer ribosomal phylogenies including previously available *Meringosphaera* sequences (Fig. 1A and *SI Appendix*, Fig. S1 and Dataset S1). The 28S rDNA tree, which received higher support values than the 18S rDNA tree for the main clades, served to delineate the main *Meringosphaera* groups (Fig. 1A). All new SAGs belonged to the formerly proposed group-2, but the diversity within this group was greatly expanded. We identified six main clades, designated MER-1 to MER-6, retaining the original numbering for group-1 for consistency while redefining the former group-2 as MER-2 through MER-5. The large MER-2 clade was further subdivided into five subclades (2A–2E).

**Figure 1.**
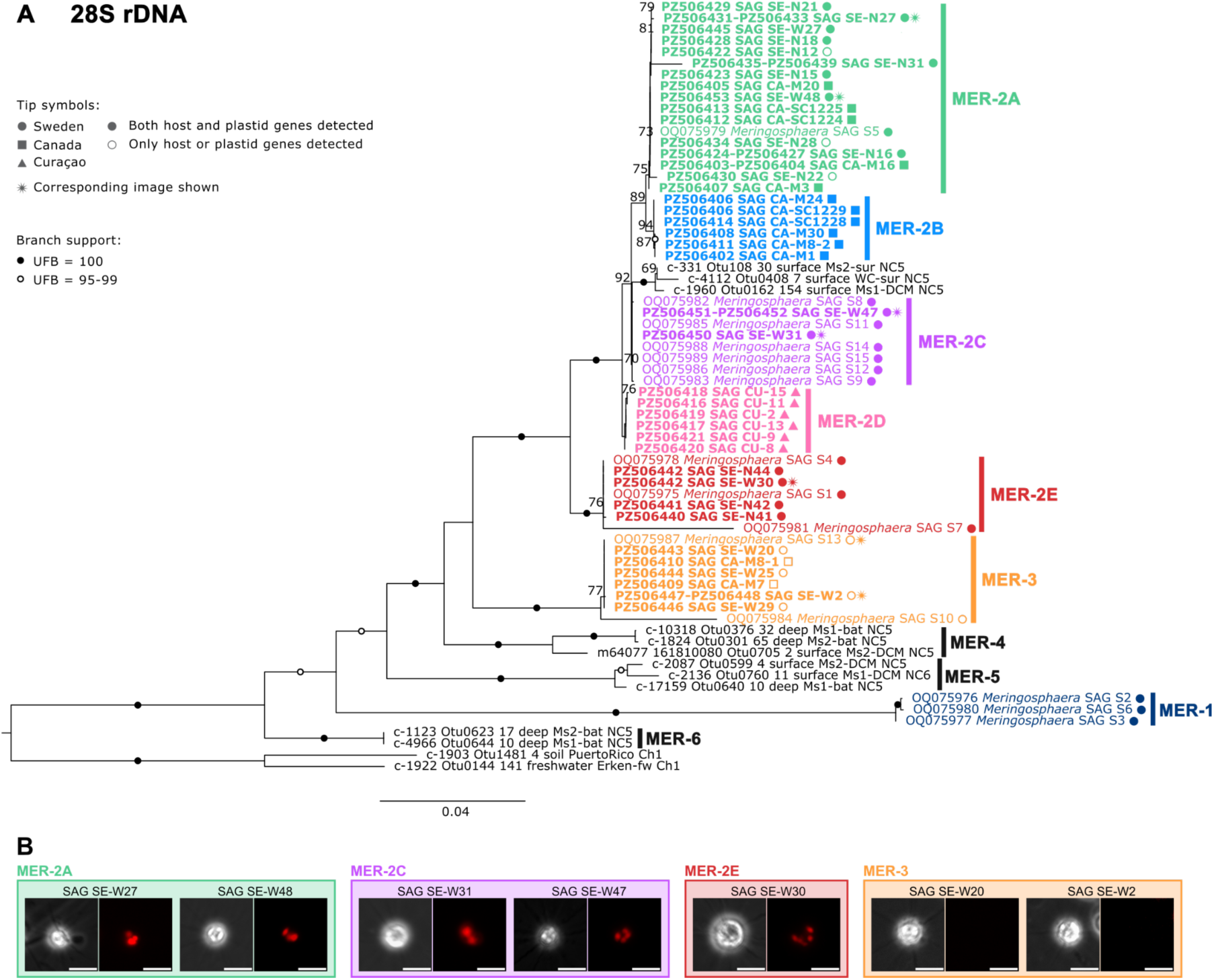
*Meringosphaera* 28S rDNA phylogeny and epifluorescent microscopy. (A) 28S rDNA phylogenetic tree depicting MER-1-6. MER-2 is further split into 2A-E. The colored clades contain SAG data. Support was calculated with 1000 ultrafast bootstrap replicates and only values equal to or greater than 70 are shown. The scale bar indicates the number of substitutions per site. Newly generated SAGs are in bold. (B) Images of *Meringosphaera* cells before isolation taken with phase contrast (left) and epifluorescence using the cy5 filter (right). Shown are two cells from MER-3 (SE-W20 and SE-W2) with no plastid autofluorescence and five cells from MER-2 (SE-W30, SE-W31, SE-W47, SE-W27, SE-W48) with plastid autofluorescence. Scale bars: 10 μm.

Currently, MER-1, MER-3, and all MER-2 subclades are represented by single-amplified genomes (SAGs), whereas MER-4, MER-5, and MER-6 comprise only environmental rDNA sequences. MER-6 is provisionally classified within *Meringosphaera*, though its status as the earliest-diverging group (Fig. 1A) may warrant future reassessment.

Contamination is a common issue with MDA, as even traces of DNA get amplified (48, 49). Contamination refers to foreign DNA accidentally introduced in the experiment, but also DNA that was present in or around the target cells during isolation, including potential prey. In addition to bacterial contamination, ten SAGs contained 18S or 28S rDNA sequences other than those of *Meringosphaera* and from various eukaryotic sources (dinoflagellate, haptophyte, stramenopile, choanoflagellate, animal, and fungi). However, we did not detect dictyochophyte nuclear-encoded ribosomal RNAs, in line with previous findings that *Meringosphaera* only contains dictyochophyte plastids but no nucleus (37). All SAGs were cleaned of major detectable contamination using a workflow combining GC content and blast similarity to a taxonomically diverse database including all available datasets for centrohelids (see Material and Methods). Post-decontamination, SAG sizes ranged from 3.5–230 Mbp, with N50 values of 859–9840 bp and GC content of 43–59% (Dataset S1). Busco completeness scores ranged from 1.5 to 58.9%, altogether reflecting a large quality disparity (Dataset S1).

### MER-2 harbors a broad diversity of plastid groups

Plastid genomes were assembled to investigate the presence, identity, and coding capacity of plastids within the new SAGs. In total, we recovered dictyochophyte plastid genome data from 38 SAGs, all belonging to MER-2 (*SI Appendix,* Fig. S2). Five genomes mapped as circular molecules, while the rest (n=33) were partial and often fragmented contigs. We compared these five new circular genomes with the previously obtained *Meringosphaera* and dictyochophyte plastid genomes (37, 50). The new plastid genomes corresponded to MER-2A, 2B, 2C, and 2E. As no plastid genome was circularised for MER-2D, we also included a partial plastid genome from this clade in our analysis (CU-15, consisting of 3 contigs amounting to 87543 bp) to cover all MER-2 groups. The plastid genome sizes of MER-2 ranged between 88-92 kbp, which is smaller than the known dictyochophyteplastid genomes (108-140 kbp) (50) (*SI Appendix,* Fig. S2 and S3). The largest plastid genome (92kbp) was observed in MER-2E, corresponding to the earliest diverging group of clade 2 (Fig. 1A). Confirming previous observations (37), all MER-2 plastid genomes encoded a highly similar gene repertoire, including the core components of photosynthesis and carbon fixation genes, with differences mainly in open reading frames (ORFs) (*SI Appendix,* Fig. S3A). All plastid genomes were highly syntenic, with only one inversion detected in MER-2E as compared to MER-2A-D, and one in MER-2A-B as compared to MER-2C-E (*SI Appendix,* Fig. S3B).

12 SAGs were without plastid data (*SI Appendix*, Fig. 2B). Of these, only three belonged to MER-2 (3/40=7.5%) but all six SAGs from MER-3 lacked plastid data. The remaining three SAGs without plastid data were of unclear phylogenetic origin as they also lacked 18S or 28S rDNA sequence, indicative of poor overall quality. SAGs are notoriously incomplete (51), and our BUSCO estimation of genome completeness is expectedly low to moderate, reaching at most 59% for SAGs without clear contamination (Dataset S1). Thus, missing plastid genomes could be due to incompleteness of SAGs. However, the complete lack of plastid sequences in not only the six new SAGs added to MER-3, but also in the two already available SAGs in this clade (SAG S10 and S13 (37)), suggest that this group is likely plastid-lacking. Supporting this hypothesis, epifluorescence microscopy of a few cells prior to MDA did not reveal chlorophyll autofluorescence in the MER-3 cells (SE-W2 and SE-W20), in contrast to all observed MER-2 cells (SE-W30, SE-W31, SE-W47, SE-W48, and SE-W27) (Fig. 1B and Dataset S1). These findings collectively indicate that MER-2 harbors plastids, whereas MER-3 does not. As no genomic or microscopic data is available for MER-4, MER-5, and MER-6, it is currently not possible to know whether these other *Meringosphaera* clades might harbor plastids.

**Figure 2.**
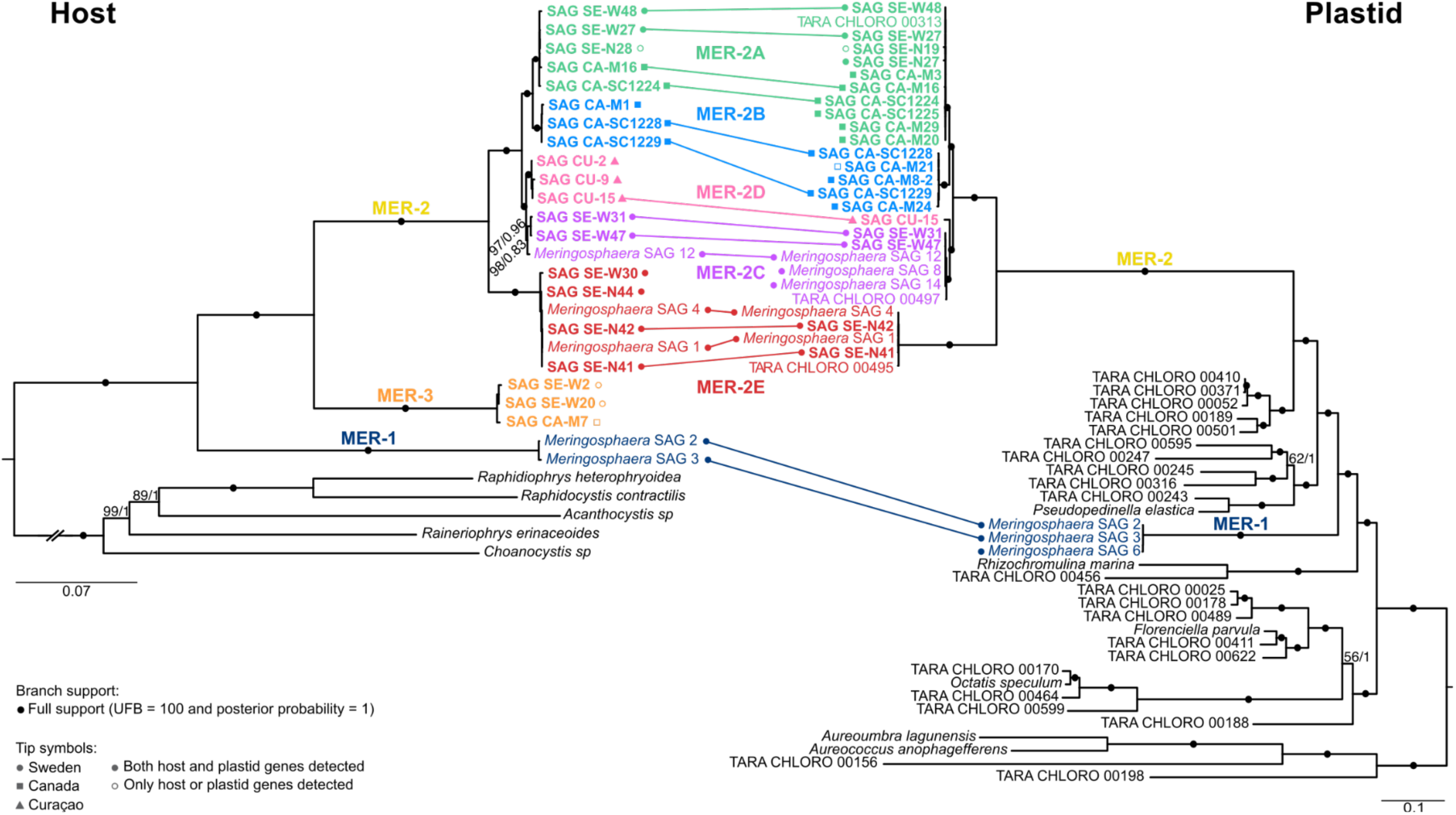
*Meringosphaera* host and plastid diversity and co-evolution. Phylogenomic trees of 88 Centrohelid nuclear genes (left) and 79 dictyochophyte plastid genes (right). The new SAGs are in bold. Support is shown as a black circle when maximal, i.e. ultrafast bootstrap support=100 and posterior probability=1. Support is shown in numbers otherwise (UFB/posterior probability). Support values within MER-1, MER-3, and MER-2A, MER-2B, MER-2C, MER-2D, and MER-2E are not shown for clarity. Host and plastid tip symbols are connected by a line when they were derived from the same SAG. The scale bar indicates the number of substitutions per site.

### Co-evolution of hosts and plastids in *Meringosphaera* clades 2A-E across geographic locations

To refine the evolutionary relationships among *Meringosphaera* host and plastid groups, we constructed phylogenomic trees (Fig. 2) using concatenated alignments of 88 host genes (with centrohelid outgroups) and 79 plastid genes (with ochrophyte outgroups). To minimize contamination risks in these multigene analyses, we restricted our dataset to genes in which MER-2A–2E formed monophyletic groups. The resulting trees, inferred with site-heterogeneous models in both Maximum Likelihood (ML) and Bayesian methods, recovered corresponding MER-2A-E clades in host and plastid in a perfectly mirroring and fully supported branching pattern (Fig. 2). Notably, MER-2A, MER-2C, and MER-2E each contained representatives from multiple locations, an observation not only based on SAG origin but also on the occurrence of environmental plastid genomes (ptMAGs) retrieved from different oceans (52) (Fig. 2 and *SI Appendix*, Fig. S4). Thus, the genetic correspondence between MER-2 host and plastid subclades does not appear to be driven by geography. In addition, the putatively plastid-lacking MER-3 group robustly branched as sister to MER-2 in the host tree. Finally, MER-1 plastids are clearly of separate origin, albeit also derived from dictyochophytes, confirming a previous inference (37).

Mirroring host and plastid phylogenies are strong indications of co-evolution. Co-evolving host-symbionts are well established in host-parasite systems and other vertically transmitted endosymbioses (53–58), but to our knowledge have not been documented in temporary relationships such as kleptoplastidy. In principle, kleptoplastidy could generate congruent host–plastid patterns, as previously proposed (37), but the expanded diversity recovered here would require repeated and highly specific associations between particular *Meringosphaera* lineages and matching free-living prey sources (Fig. 3A). Instead, we propose that MER-2 possesses vertically transmitted permanent plastids acquired from dictyochophytes in an ancestor of the group, likely after its divergence from MER-3, followed by diversification into at least five subclades (Fig. 3B). MER-1 may also possess permanent plastids, but more data is required to fully evaluate the endosymbionts in this group. Notably, neither MER-1 nor MER-2 plastids branch closely to any currently sampled dictyochophyte lineage, even after inclusion of ptMAGs in addition to the four named species with available plastid genomes. If these organelles were kleptoplasts, they would therefore have to derive from an unsampled prey source. In this new context, the previous interpretation of kleptoplastidy in MER-2 (37) is best understood as a consequence of limited taxon sampling. The addition of SAGs from multiple regions and seasons revealed previously unrecognized diversity (especially within MER-2A) that was not captured by the earlier plastid-targeted qPCR assay. As a result, patterns previously interpreted as plastid switching are more parsimoniously explained by stable associations between distinct host lineages and their own vertically inherited plastids.

**Figure 3.**
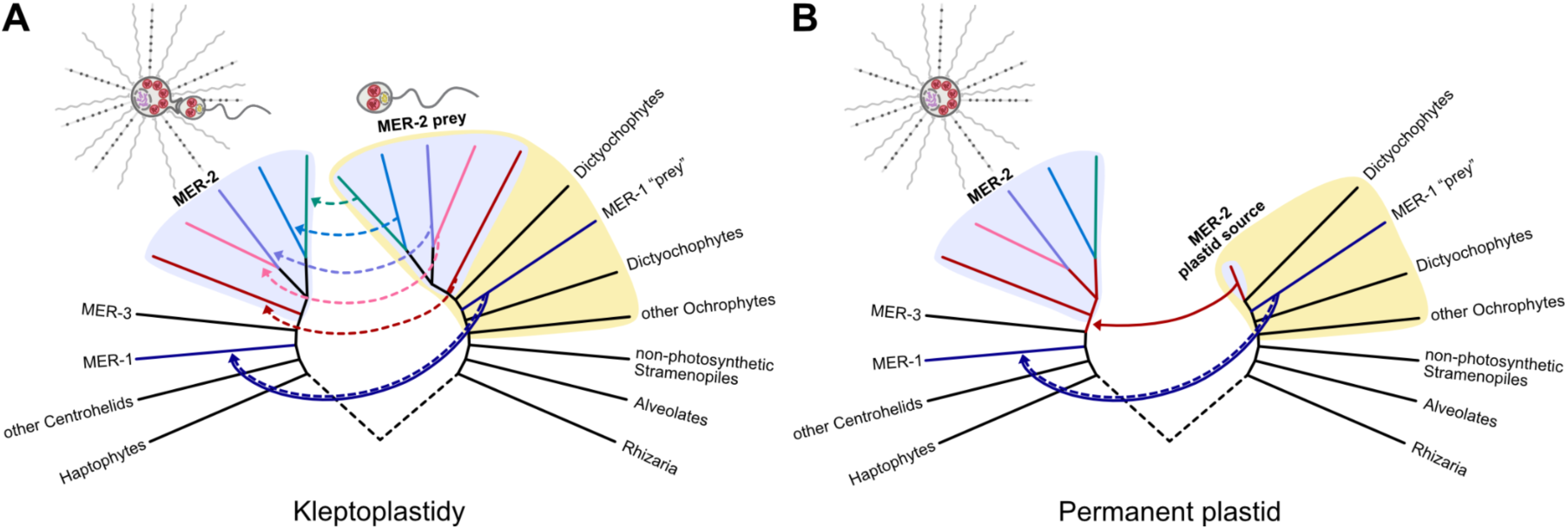
Kleptoplastidy versus permanent plastid hypotheses in MER-2. Illustrations of part of the eukaryotic tree of life under the hypothesis that MER-2 performs kleptoplastidy (A) or harbors a permanent plastid (B). Solid arrows indicate permanent integrations and dashed arrows indicate kleptoplastidy. MER-2 host and plastid are highlighted in purple and ochrophytes are highlighted in yellow. As it is currently unclear whether MER-1 harbors permanent plastids or kleptoplastids, a combined solid and dashed arrow is shown for this group.

### Double CARD-FISH shows high prevalence of plastids in *Meringosphaera* clade 2

Using the extended diversity of MER-2, we established a double CARD-FISH assay to simultaneously target the MER-2 subclades for both host (18S rRNA) and plastid (16S rRNA). The rationale was two-fold: (1) to investigate more systematically than with SAGs the prevalence of MER-2 plastids in environmental samples, and (2) to search for hypothetical cells displaying a plastid signal but no host signal, which could correspond to plastids in potential prey. A total of 119 *Meringosphaera* cells were detected and imaged from 19 filters collected at the same locations as the SAGs (27 from Canada, 43 from Curaçao, and 49 from Sweden) (Fig. 4 and Dataset S2). Combined with the 50 new SAGs, this study thus encompasses 169 environmental *Meringosphaera* cells. Each cell was evaluated for probe hybridization strength and localization, chlorophyll autofluorescence, and the presence of a DAPI-stained nucleus. We also assessed whether the cells displayed the typical *Meringosphaera* morphology with a round body of the expected size range (3-12 um in diameter). In many cases, the characteristic undulating spines were also observed. The absence of spines could result from sample handling and processing, as *Meringosphaera*’s spines were easily dislodged and were sometimes observed close, but not attached, to the *Meringosphaera* cell body, or in other places on the filter.

**Figure 4.**
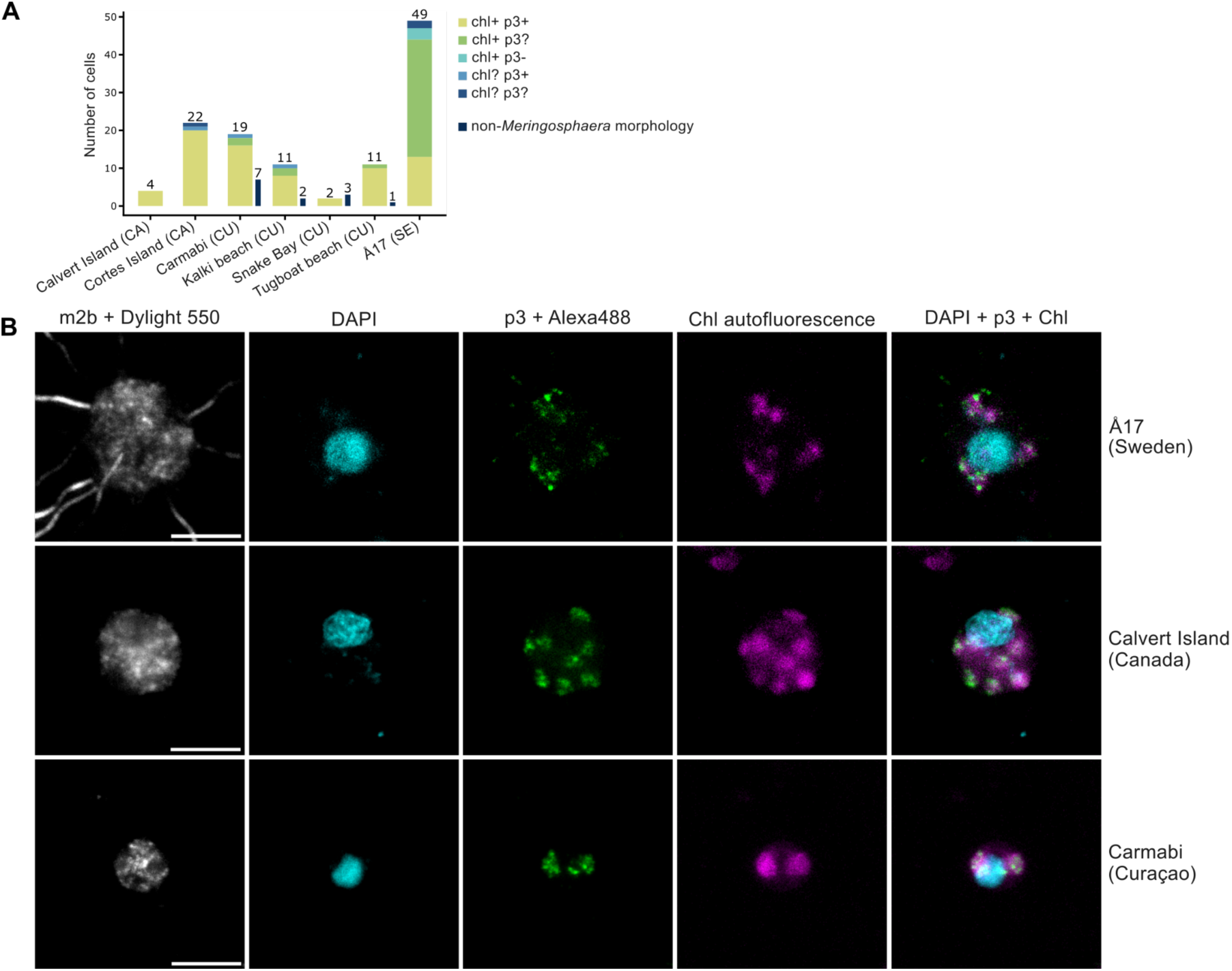
Double CARD-FISH of *Meringosphaera* and cells with non-*Meringosphaera* morphology but positive plastid probe hybridization on samples from Sweden, Curaçao, and Canada. (A) A Bar graph showing the number of *Meringosphaera* cells as well as the number of cells with non-*Meringosphaera* morphology imaged from samples from each location. Legend shows the different colors represent different levels of positive (+), uncertain (?) and negative (−) chlorophyll autofluorescence and plastid 16S rRNA probe (p3) signal. (B) Three *Meringosphaera* cells from samples from Å17 (Sweden), Calvert Island (Canada), and Carmabi (Curaçao). From left to right is shown the host 18S rRNA probe (m2b) with amplified Dylight550 fluorophore, DAPI signal, plastid 16S rRNA probe (p3) with amplified Alexa488 fluorophore, chlorophyll autofluorescence imaged with the 633 laser, and an overlay of the DAPI, p3+Alexa488, and chlorophyll autofluorescence signals. Scale bar is 5 μm.

Of the 119 imaged *Meringosphaera* cells, 116 exhibited visible plastids (2–12 when enumerable), as indicated by clear chlorophyll autofluorescence, positive hybridization with the MER-2 plastid probe, or both (Fig. 4A and Dataset S2). The absence of chlorophyll autofluorescence in some cells likely resulted from bleaching during CARD-FISH chemical incubation (59–61), particularly as 16 of these cells showed positive MER-2 plastid probe hybridization. Additionally, 39 cells (primarily from Sweden) displayed chlorophyll autofluorescence but ambiguous MER-2 plastid probe hybridization due to weak or undetectable signals against the background. Two main explanations can account for the poorer signal of the plastid probe with the Swedish samples.

First, it is possible that some cells carried a different plastid unspecific to the probe. However, we consider this possibility unlikely due to the very high specificity we observed between the *Meringosphaera* hosts and their plastid across clades 2A-E. Alternatively, it is possible that the *Meringosphaera* cells had a short lifespan after being sampled and the plastids started to degrade rapidly, which could be reflected in lower 16S rRNA content of the plastids, leading to weak signal of the 16S rRNA probe. Unlike for other locations, the Swedish samples were chemically fixed with a 3-4 days delay after collection, thus possibly triggering degradation.

In addition to these 119 cells, we also found 13 plastid-positive cells in samples from Curaçao that did not have the characteristic *Meringosphaera* morphology (Fig. 4A and *SI Appendix* Fig. S5 and Dataset S2). These cells were smaller (2-4 μm in diameter, including smaller nucleus), elongated, and consistently had two plastids flanking the nucleus (*SI Appendix* Fig. S5 and Dataset S2). It is possible that these cells represent an unknown lifestage of *Meringosphaera*, as some centrohelids are known to demonstrate dimorphism, including dramatic size and morphology differences (62). Alternatively, the positive probe signal could derive from unspecific targeting to other unknown algae with similar plastid rRNA sequences, or correspond to a dictyochophyte prey source of hypothetical kleptoplasts. However, given our phylogenomic inference and overall CARD-FISH signal indicating a permanent plastid in MER-2, we consider the latter explanation less plausible. Furthermore, we would also expect to detect these cells in samples from Canada and Sweden, but none were found.

### *Meringosphaera SAG* co-assemblies reveal ancestral acquisition and diverse origin of nuclear-encoded core plastid metabolism pathways

To assess the extent of plastid integration into the host genome, we searched *Meringosphaera* coassemblies (COSAGs) for 70 host-encoded genes typically associated with plastid function in algae. These included genes involved in core plastid metabolism such as fatty acid biosynthesis (5 genes), heme biosynthesis (7 genes), isoprenoid biosynthesis (7 genes), and iron–sulfur cluster assembly (7 genes), as well as plastid protein import (34 genes), plastid division (8 genes), the chaperone protein *cpn60*, and the nuclear-encoded photosynthetic gene *psbO*, which encodes the manganese-stabilizing protein of Photosystem II. To improve genome recovery, we coassembled only SAGs sharing highly similar 18S rDNA sequences (>99% identity) which were assigned the same subclade and originated from the same country, yielding nine COSAGs spanning all MER-2A–E clades, MER-3, and previously available MER-1. BUSCO completeness ranged from 18% to 70%, representing an average improvement of 11% over individual SAGs (Fig. 5A, Dataset S1 and *SI appendix* Table S1). We then used phylogenetic analyses with broad taxonomic sampling (Dataset S3) to identify *Meringosphaera* homologs and exclude likely contaminants. Additional centrohelid datasets were essential for distinguishing plastid-associated homologs encoded by *Meringosphaera*, which grouped with algae, from mitochondrial or cytosolic homologs, which grouped with centrohelids (Fig. 5A). Whenever possible, we required consistent placement across multiple COSAGs to support gene assignment. The only exceptions were *psbO* and *dxr*, each recovered from a single COSAG but present in two individual SAGs used in the coassemblies.

**Figure 5.**
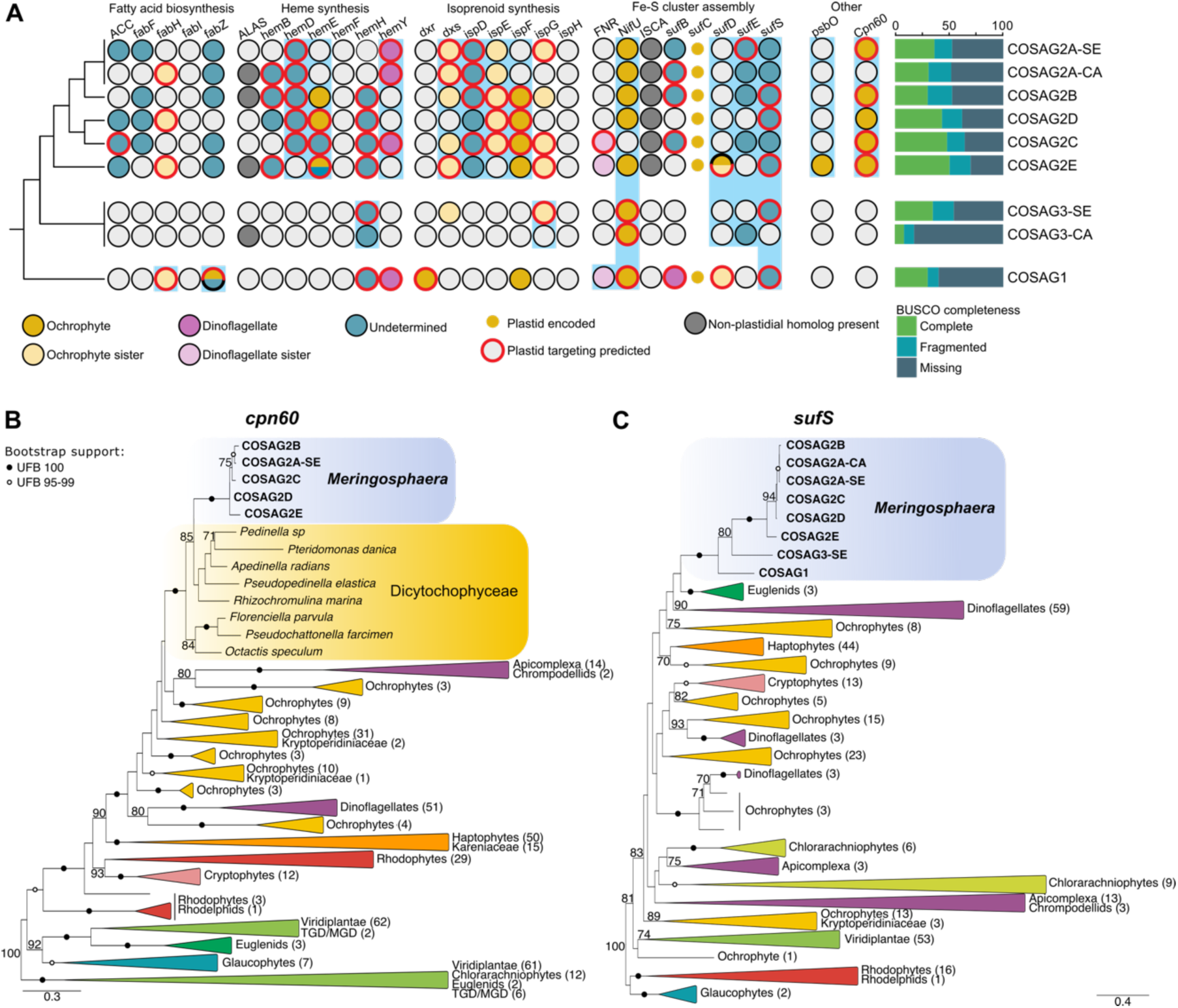
Host-encoded plastid associated genes in nine *Meringosphaera* co-assemblies and their phylogenetic origin. (A) The presence, phylogenetic origin, and predicted localization of plastid-associated genes encoded by nine *Meringosphaera* COSAGs. A blue box is present behind the genes that were not previously known to be present in MER-1, MER-2, or MER-3. Busco completeness scores of each COSAG are shown on the right. (B and C) Phylogenetic tree of *cpn60* and *sufS*, showing the phylogenetic placement of these host-encoded *Meringosphaera* genes. Support was calculated with a thousand ultrafast bootstrap replicates and only shown on the tree if it was higher than 70. Support of 100 UFB is shown as filled black circles and 95-99 UFB is shown as empty circles.

Overall, we identified 20 nuclear-encoded genes involved in core plastid metabolism in MER-2, providing strong evidence for genetic integration in this lineage (Fig. 5A and Dataset S4). Their distribution was patchy, likely reflecting incomplete genome recovery. Some genes, such as *ispD* and *sufS*, were present in all MER-2 COSAGs, whereas others, including *fabI*, *hemF*, *ispH*, and *dxr*, were not detected. Although these absences may reflect MDA bias, some genes may also be genuinely missing. MER-2 also encoded *psbO*. Although many photosynthesis-related genes are plastid-encoded in algae, and many are retained in the *Meringosphaera* plastid genome (*SI Appendix,* Fig. S3), *psbO* is consistently nuclear-encoded (63, 64). Its presence therefore suggests host control over part of the photosynthetic machinery. We also detected plastid *cpn60* in five MER-2 COSAGs. This chaperone has key plastid functions, including folding and assembly of imported proteins, promotion of Rubisco assembly, and regulation of plastid division (65–68).

In MER-1, which also harbors plastid genomes, we detected 11 nuclear-encoded genes involved in core plastid metabolism in the single available COSAG, but not *psbO* or *cpn60*. In MER-3, which likely lacks a fully integrated plastid (see above), we detected fewer plastid-related genes (7 genes) and also didn’t find *psbO* or *cpn60*.

Because plastids depend on the import of nucleus-encoded proteins, the evolution of targeting and import systems is generally viewed as a key step in permanent organelle establishment (16, 32). Plastid-targeting sequences were predicted in many plastid-associated proteins in MER-2 (17/22) and MER-1 (9/11), but also in MER-3 (4/5). In several MER-2 genes (*fabH*, *hemH*, and *ispD*) and one MER-3 gene (*sufS*), we detected a potential bipartite targeting signal consisting of a signal peptide followed by a chloroplast transit peptide within 100 bp, a pattern often associated with complex plastids (Dataset S4, *SI Appendix,* Fig. S6) (69). In other cases, only one motif was detected, or the two occurred in reversed order or in different positions. Although *cpn60* also occurs in mitochondria, the MER-2 copies are clearly of dictyochophyte origin (Fig. 5B), and four out of the five detected copies (across five MER-2 COSAGs) contain an N-terminal plastid-targeting signal, supporting plastid localization.

Despite the evidence for targeting, we found no clear homologs of canonical plastid import machinery encoded in the *Meringosphaera* host genome (see Material and Method for a list of genes). We detected no convincing components of the TIC/TOC machinery, which mediates transport across the two innermost plastid membranes, nor of the symbiont-derived ERAD-like machinery (SELMA), which facilitates transport across third membrane of red algal-derived plastids, including tertiary plastids (15, 16, 70, 71). We did find two of the components of the host’s Sec61 complex (the alpha and gamma components), which mediates transport across the outermost plastid membrane in several algae (69, 72). However, Sec61 is a central component of the protein translocation machinery not only in algae, but in general facilitating the co-translational transport of proteins across the endoplasmic reticulum (ER) membrane. Thus, the presence of this complex in *Meringosphaera* is expected and branched with or close to those of other centrohelids. Taken together, these results indicate that *Meringosphaera* may use a distinct import strategy, which is confounded by the fact that the number of plastid membranes remain unknown. Similar departures from canonical systems have been reported in other algae, including euglenids and *Paulinella* (35, 73–75). As for import proteins, our search for plastid division genes, including *ftsZ*, ARC5/DRP5B, and *minC–E*, yielded no convincing candidates. This is not unexpected, as plastid division systems remain poorly characterized in algae with complex plastids (76).

The phylogenetic origins of the nuclear-encoded plastid proteins were mixed. For *cpn60*, a dictyochophyte origin was strongly supported, consistent with an ancient transfer into a *Meringosphaera* ancestor (Fig. 5B). This gene most likely derived either from the endosymbiont itself or from a closely related lineage. In 11 additional cases, the best-supported origins lay within ochrophytes other than dictyochophytes (for example, diatoms, bolidophytes, and/or pelagophytes) or more broadly within unresolved ochrophytes. These genes may also derive from the endosymbiont, although alternative donor lineages cannot be excluded. Two *Meringosphaera* genes showed affinity to dinoflagellates. For most genes, the origin remained unresolved, either because *Meringosphaera* branched outside defined algal clades or because deeper relationships lacked support (Fig. 5A). Notably, in 12 of the 13 cases in which homologs were detected in MER-2 as well as MER-1 and/or MER-3, these different *Meringosphaera* clades grouped together (Fig. 5C). This pattern indicates that at least some plastid-related genes were acquired before the divergence of MER-1, MER-2, and MER-3, and therefore before plastid integration.

In summary, these results indicate that *Meringosphaera* has not only incorporated foreign algal genes into its nuclear genome but also evolved at least partial host control over plastid metabolism. The mixed phylogenetic origins of these genes, the acquisition of some before organelle fixation, and the presence of putative plastid-targeting signals even in MER-3 are all consistent with predictions of the shopping bag model of plastid origin (31, 32). Although this model has previously been supported in kareniacean dinoflagellates and the kleptoplastidic euglenid *R. viridis*, *Meringosphaera* extends it to a phylogenetically distinct system unrelated to known algal groups, suggesting that these processes may reflect broadly applicable principles of plastid evolution.

## Conclusions

*Meringosphaera* is an uncultured group of marine centrohelids that harbors dictyochophyte plastids from at least two independent origins. Based on a narrow sampling, we previously suggested that these organelles were kleptoplasts stolen from an unidentified dictyochophyte prey. Here, based on a combination of phylogenomics, fluorescent microscopy, and comparative genomics encompassing many more samples and representing a much wider range of diversity, it is now clear that in at least one subgroup (MER-2) the plastids are permanently integrated.

Historically, *Meringosphaera* has often been (but not always) classified as an algal group based on sporadic microscopic observations. Our results fit well with the observation that superficially similar cells can belong to both plastid-bearing and plastid-lacking groups. Indeed, without epifluorescence microscopy, MER-2 and MER-3 appear morphologically indistinguishable.

Moreover, our findings provide strong evidence for algae in centrohelids. Novel algae not part of already established algal groups are very rarely discovered, with only two examples in the last 100 years (52, 77, 78). Thus, the report of centrohelid algae demonstrates the existence of cryptic photosynthetic diversity that further complicates an already very convoluted evolution of plastid endosymbioses. Yet, *Meringosphaera* also helps better understand the early steps of plastid establishment and, in the absence of a culture, shows the importance of studying environmental cells with culture-free approaches.

## Materials and Methods

### Sample acquisition

Samples were collected between surface water to 30m deep from the station Å17 on the west coast of Sweden by the Swedish Meteorological and Hydrological Institute (SMHI) monthly between May 2022 and October 2023, from Calvert Island, Cortes Island, Ogden Point, and Quadra Island on the Pacific Northwest coast of Canada between June and July of 2024, and in front of the Carmabi research station as well as at Kalki beach, Tugboat beach, and Snake Bay in Curaçao in October of 2024 (Dataset S1 and S2). In most cases whole water samples were collected by bottle or bucket, but some samples were taken by gentle plankton tows (Calvert Island, Quadra Island, and some samples collected in Cortes Island). Water was processed between 0 to 14 days after collection and when transport and storage was needed, it was cooled with ice packs or kept in a cold room in the dark.

### Single cell isolations and MDA

Single cell isolation and multiple displacement amplification (MDA) of *Meringosphaera* cells were performed similarly as described in (37). Briefly, after receiving the whole water samples, approximately 500 ml was concentrated using gravity onto a 47 mm diameter 5 μm pore size Merck Isopore or Cytiva Whatman Cyclopore polycarbonate membrane filter. Cells were resuspended from the filter in 10 ml of filtered seawater (FSW) for approximately one minute.

Plankton tow samples were not further concentrated after sampling. *Meringosphaera* cells were visually identified with a Nikon Eclipse Ts2R-FL inverted microscope using phase contrast. The cells were imaged before isolation and, when possible, they were also assessed for chlorophyll autofluorescence signal using the Semrock Cy5-4040C filter set (excitation: 628/40-25, emission: 692/40, dichroic: FF660-Di02). Single cells were isolated either using an Eppendorf TransferMan 4r micromanipulator with pulled glass capillaries, or picked by hand using pulled glass capillary attached to a rubber tube connected to a syringe. The cells were washed in UV-sterilized 0.2 μm FSW 0-3 times and then inserted into a UV-sterilized PCR tube, before being frozen at −20 or −70 °C until MDA. MDA was performed with either the REPLI-g UltraFast Mini kit (Qiagen) or the REPLI-g Advanced DNA Single Cell kit (Qiagen) (see Dataset S5 for SAG specification). After amplification, MDA products were screened for the presence of *Meringosphaera* 18S rDNA with the primers Thx25F (5’-CATATGCTTGTCTCAAAGATTAAGCCA-3’) and Helio1979R (5’- CACACTTACWAGGAYTTCCTCGTTSAAGACG-3’) (79). The MDA products with correct-sized bands were sent for Sanger sequencing at Macrogen (Netherlands) to confirm the presence of *Meringosphaera*.

### Sequencing

50 MDA products with confirmed *Meringosphaera* 18S rDNA sequence were sent for sequencing at SciLifeLab National Genomics Infrastructure (NGI), Sweden. For *Meringosphaera* cells collected between May and August of 2022, libraries were prepared using the TruSeq PCR-free kit, except with the TruSeq Nano kit in one case (SAG SE-N44) due to limiting DNA concentration, targeting an insert size of 2×150bp (Dataset S5). Samples were multiplexed on 2 lanes and sequenced on a NovaSeq 6000 (Illumina) with pair-end standard setup and a SP-300 v1.5 flowcell. For *Meringosphaera* cells collected after August 2022, libraries were prepared using the NGI in-house Splinted Ligation Adapter Tagging (SPLAT) protocol excluding sodium bisulfite conversion (80) (Dataset S5). Samples were multiplexed on a 10B flowcell using XLEAP-SBS sequencing chemistry with a NovaSeq X Plus system (Illumina), targeting a paired-end 150bp read length.

### SAG assembly and phylogenetic analysis of the rDNA

All reads were first trimmed with TrimGalore v0.6.1 (https://www.bioinformatics.babraham.ac.uk/projects/trim_galore/) and subsequently normalized to a target value of 25 with bbmap v.38.08 (81) or to a target value of 100 with bbmap v.39.08 (Dataset S5). The normalized reads were assembled into contigs with Spades (82) v.3.15.5 with metaspades.py or Spades v.4.0.0 with spades.py using the option --careful (Dataset S5).

Different normalization values and assembly strategies were first tested on a subset of the data and the quality of the resulting assembly tested using quast v.4.5.4 or v.5.0.2 and busco v.5.7.1, before choosing the best strategy. The 50 resulting Single-cell Assembled Genomes (SAGs) were searched for 18S rDNA and 28S rDNA sequences using Barrnap v.0.9 (https://github.com/tseemann/barrnap). For phylogenetic analysis, the *Meringosphaera* 18S rDNA and 28S rDNA sequences were aligned with representative outgroup centrohelid sequences obtained from Genbank, PR2, and the long-read OTUs from (83) with MAFFT (84) G-INS-i v.7.526 and trimmed with TrimAl (85) v.1.5.0 (-gt 0.1). Trees were constructed with IQ-TREE (86) v.3.0.1 using ModelFinder (87) to select the best model and support values were calculated with a 1000 ultrafast bootstrap replicates (-m MFP, -B 1000). The best-fit model for both the 18S rDNA and 28S rDNA phylogenetic trees was TN+F+R3.

### Plastid genome assembly and comparisons

Plastids were assembled using GetOrganelle (88) v.1.7.7.0 from reads normalized with bbmap (81) v.39.08 to a target value of 100 using previously assembled *Meringosphaera* plastids from MER-2 as a seed (-F other_pt -R 15 -k 21,45,65,85,105). The plastid genomes were annotated with MFannot (https://megasun.bch.umontreal.ca/apps/mfannot/) and the content was checked with OGdraw (https://chlorobox.mpimp-golm.mpg.de/OGDraw.html). The gene annotation was manually corrected. Subsequently, OrthoFinder v.2.5.5 was used on the protein files, choosing the correctly rooted tree (dictyochophyte outgroup), to assess their annotation. Single gene trees generated by OrthoFinder were manually inspected and the misannotation of *rpl22* as simply orf119 was corrected. An R script from (52) was used to visualize the orthofinder results. Plastid genome synteny analysis was performed with pyGenomeViz v1.6.1 (https://pygenomeviz.streamlit.app/) using the MMseqs RBH genome comparison method.

### Phylogenomic analyses

#### Host data

We generated predicted proteomes of the new *Meringosphaera* SAGs alongside the 15 previously published *Meringosphaera* SAGs (37) using Prodigal (89) v.2.6.3 or MetaEuk (90) easy-predict using the Tara Oceans protein database published along with MetaEuk as a reference (Dataset S5). From these predicted proteomes we collected homologs of 317 well-conserved proteins using the PhyloFisher (91, 92) pipeline in two consecutive rounds (Dataset S5). Briefly, we collected *Meringosphaera* homologs and combined these with the PhyloFisher dataset using fisher.py and working_dataset_constructure.py and subsequently constructed single gene trees manually. We used PREQUAL (93) v.3 on each collected *Meringosphaera* homolog separately to mask intron regions and non-homologous residues, followed by alignment with MAFFT (84) G-INS-i v.7.407 unalign level 0.6 and light trimming with TrimAl (85) v.1.4.1 (-gt 0.1). We then constructed individual gene trees with IQ-TREE (94) v.2.2.2.6 using the model LG4X+G and calculating support values with optimized ultrafast bootstraps (-B 1000, -bnni). Initial assessment of the resulting alignments and single gene trees revealed that many *Meringosphaera* sequences were fragmented. We manually merged these sequence fragments from the same proteome only if (1) they overlapped and all overlapping amino acids were the same, (2) non-overlapping fragments were encoded sequentially on the same contig, (3) the fragments branched as direct sisters, or (4) their relationship was unresolved but multiple *Meringosphaera* proteomes showed the same pattern of sequence fragmentation. After this sequence fragment merging, homologs were re-aligned, trimmed and single gene trees constructed as described before. All single gene trees were manually parsed to determine orthology and identify contaminants, and these decisions were applied to the PhyloFisher (91, 92) dataset using apply_to_db.py. We constructed phylogenomic trees including only proteins that were present in at least one representative proteome of clade 2A-E, and only taxa that had at least 20% of the selected proteins. Protein sequences were aligned with MAFFT (84) G-INS-i v.7.407 (--unalignlevel 0.6) and trimmed with BMGE (95) v1.12 BLOSUM35 matrix, gap threshold: 0.8, before concatenation of the alignments. Phylogenomic trees were generated with IQ-TREE (94) v.2.2.2.6 using ModelFinder (87) to find the best fitting LG+C60 site-heterogeneous model (LG+C60+F+I+R3). Support was calculated with optimized ultrafast bootstraps (-B 1000, - bnni). Finally, we also built a phylogenomic tree using PhyloBayes (96) with the CAT-GTR model (2 chains for 2700 cycles and burning of 500).

#### Plastid data

We first extracted plastid contigs with BLAST (97) v.2.15.0+ using previously published *Meringosphaera* and dictyochophyte plastid genomes and plastid metagenomes as queries (37, 50, 52). Proteome prediction of the extracted plastid contigs as well as of the previously published *Meringosphaera* plastid genomes (37) was performed with Prodigal (89) v.2.6.3. We then used the plastid protein database generated in (52) including 93 well curated plastid proteins from all major algal lineages as well as plastid metagenomes assembled from Tara Oceans data. As above, we performed our analysis in two consecutive rounds (Dataset S5). These 93 proteins were used as queries for BLASTp v.2.15.0+ against the plastid proteomes to collect *Meringosphaera* homologs, and hits with at least 0.5 protein coverage and an e-value of 1e-20 were retained and combined with the initial dataset. Alignment, trimming, single gene tree construction and sequence fragment merging was performed as above. All resulting trees were parsed to remove contaminants and paralogs. Similar to our strategy for the host phylogenomic analysis, we only included plastid proteins if they were found in at least one representative of clade 2A-E and taxa that had at least 50% of these selected genes. We aligned all proteins individually with MAFFT (84) G-INS-i v.7.407 (--unalignlevel 0.6), trimmed them with BMGE (95) v1.12 BLOSUM35 matrix, gap threshold: 0.8, and then concatenated the alignments. We then built a phylogenomic tree with IQ-TREE (94) v.2.2.2.6 with the best fitting cpREV+C60 site-heterogeneous model (cpREV+C60+F+I+R5) using modelfinder (87) (-MFP) and computed support with a 1000 optimized ultrafast bootstraps. We also constructed a tree using PhyloBayes (96) with the same parameters as for the host.

### Double CARD-FISH

Between 0.4 to 1 liter of water was filtered first through a 5 µm filter by gentle gravity filtration and then through a 1 or 0.2 µm filter with a peristaltic pump or vacuum pump. Filters were fixed with 4% paraformaldehyde (PFA) or 3.7% formaldehyde (FA) before the filters dried out and left in the fixative for 45-90 minutes (up to 3 hours). Separate samples were fixed before filtration at a volume of 90 ml seawater and 10 ml 37% FA (for a final concentration of 3.7% FA) for at least an hour and then filtered through a 1 or 0.2 µm filter (See Dataset S2 for specification per filter). In both cases, the fixative was rinsed off with 0.2 µm filtered seawater three times and filters were air-dried and subsequently stored at −20 degrees.

Double CARD-FISH probes were designed using the Oligo-N probe design pipeline (98), followed by additional checks of probe specificity against the PR2 and SILVA databases (99, 100) as well as checks for homo-dimers and hairpins using the IDTDNA OligoAnalyzer tool. Probes were also aligned back against the target sequences to further check specificity and mismatches. Two probes were designed targeting MER-2: one targeting the 18S rRNA (Mer2b_N6572 (m2b for short), 5’-TGCCGATCAACGGTCAGACTACG-3’) and one targeting the plastid 16S rRNA (MerChp935 (p3 for short), 5’-GTATGCCATGTCAAACCCTG-3’). Probe function was tested at a hybridization temperature of 35 °C with hybridization buffers of 20%, 30%, 40%, 50%, and 60% formamide and final probe concentration of 5, 1, and 0.5 ng/μl. Final concentrations of 1 ng/μl and hybridization buffer of 60% formamide and 0.5 ng/μl and hybridization buffer of 50% formamide were selected for Mer2b_N6572 and MerChp935, respectively.

We followed a modified version of the CARD-FISH protocol described in (101). Briefly, filters were embedded in 0.1% agarose and underwent an initial permeabilization step by incubating them in 10 mg/ml lysozyme solution for 1 hour at 37 °C. Filters were quickly washed with distilled water and moved to 0.01M HCl for a 10 min incubation at room temperature. Filter pieces were then hybridized with probe MerChp935 for approximately 20 hours at 35 °C in 40 ml hybridization buffer. Subsequently, washing was done at 37 °C for 2x 10 min with shaking in a washing buffer with appropriate NaCl concentration related to the 50% concentration of formamide in the hybridization buffer (0.019 M). Filter pieces were equilibrized in 1x PBS for 15 min at room temperature and then incubated in an amplification buffer with Alexa Fluor 488 for 1 h at 45-47 °C. Amplification buffer was washed off twice with 1x PBS for 10 min at room temperature with shaking and then the filter was quickly washed in distilled water. The MerChp935 probe was inactivated by incubating the filter in 0.01M HCl for 10 minutes, before a quick wash in distilled water and subsequent airdrying. Then, the filter was hybridized overnight with the Mer2b_N6572 probe and appropriate hybridization buffer (with 60% formamide) and respective NaCl concentration in the subsequent washing buffer (0.009 M). The second CARD-FISH proceeded like the first CARD-FISH, but with DyLight 550. After PBS washing, quick washing with distilled water, and air drying, the filter pieces were counterstained and mounted with a glycerol-based DAPI anti-fading mounting solution (final DAPI concentration: 5 μg/ml) on a glass slide and with a 1.5 thickness coverslip.

Mounted filter pieces were imaged with a CLSM Leica Stellaris confocal microscope, using the preset setting for imaging DAPI (405 nm excitation), alexa 488 for the 16S rRNA probe (498 nm excitation), alexa 546 for the 18S rRNA probe (for DyLight 550, since no setting for dylight 550 was available, 555 nm excitation), and alexa 633 to image chlorophyll autofluorescence (631 nm excitation). Tau sense was also activated to facilitate separation of probe signal and chlorophyll autofluorescence from background. Maximum intensity projection images were generated from the z-stacks with Fiji v.2.0 and all images were further analyzed with Fiji and Rstudio v.4.3.2. All cells were scored for the hybridization of the plastid 16S rRNA probe MerChp935 as well as for the presence of chlorophyll autofluorescence signal indicating plastids. Due to heterogeneity in signal intensity across the different samples, successful hybridization was assessed by 1) signal strength compared to filter background; 2) localization of the probe signal, expected to appear in distinct patches in the cell as well as overlap with the chlorophyll autofluorescence when present. Cell and nucleus diameters were calculated as equivalent circle diameters (defined as diameter = √(4*region of interest (ROI) area)/π) and all ROIs were manually selected. Statistical analysis comparing cell and nucleus diameters of *Meringosphaera* and unidentified non-*Meringosphaera* cells was performed using the Wilcoxon rank-sum test.

### SAG cleaning and Co-assemblies

All 65 *Meringosphaera* SAGs (including those previously published (37)) were cleaned with Blobtools (102) v.1.1.1 to remove contamination. Briefly, in preparation of future co-assembly, all SAGs were first assembled with Spades (82) v.4.0.0 (--careful) from reads normalized to a target value of 100 with bbmap (81) v.39.08. Additional input files for Blobtools included bam files generated with samtools (103) v.1.20 and bwa-mem2(104) v.2.2.1, and blast output files (-outfmt 6) generated by using our assemblies as queries in a blast against the genbank database (97) as well as available centrohelid genomic and transcriptomic data (105–107) and dictyochophyte transcriptomes downloaded from Eukprot (108) using BLAST (97) version 2.16.0+. After a filtering step excluding all hits with a bitscore lower than 200, there were no blast hits remaining with the dictyochophyte transcriptomes. After visualization and assessment with Blobtools (102), we removed contigs with a GC content outside of 0.4-0.8, non-eukaryotic contigs bigger than 10 kbp, and specific eukaryotic contamination per SAG (determined by 18S/28S rDNA presence in the SAGs). The normalized reads per assembly were then mapped to these “cleaned” contigs using SAMtools (103) v1.20, BWA (109) v.0.7.18, and BEDTools (110) v.2.31.1. “Cleaned” reads of SAGs were included for co-assembly based on completeness (i.e., including at least ten out of the 317 proteins searched during the host phylogenomic analysis (see above)), and low contamination level. Reads were co-assembled based on an 18S rDNA identity of at least 99%, phylogenomic (sub)clade, and country of origin using Spades (82) v.4.0.0 --careful. Co-assemblies were assessed with Blobtools like above. All contigs whose top 5 hits against Genbank were bacterial and had no hits against our local blast databases were removed. Reads were mapped back to the cleaned contigs like described previously, and co-assembled again using Spades v.4.0.0 using single cell mode (--sc) to take into account coverage differences across different SAGs generated by MDA. Afterwards, all contigs smaller than 500 bp as well as *Meringosphaera* plastid contigs were removed. Co-assemblies were assessed with Quast (111) version 5.3.0 and BUSCO (112) version 5.8.2 (*SI* appendix, Table S1) and proteomes were predicted with MetaEuk (90) with the Tara Oceans dataset as reference as described above.

### Analyses of host-encoded plastid associated genes

We searched the nine *Meringosphaera* co-assemblies alongside a local database of 421 proteomes representing all major eukaryotic groups as well as bacteria, enriched for algal lineages (especially Ochrophytes) as well as Centrohelids (Dataset S3) for the presence of core plastid metabolism pathways (5 fatty acid biosynthesis genes, 7 heme biosynthesis genes, 7 isoprenoid biosynthesis genes, and 7 iron-sulfur-cluster assembly genes, see Fig. 5A), 15 TIC/TOC components (TIC20, TIC21, TIC22, TIC32, TIC55, TIC56, TIC62, TIC100, TIC110, TIC214, TOC33, TOC34, TOC64, TOC75, TOC159), 16 SELMA components (Der1, Dfm1, PUBL, Uba1, Ubc, Hrd1, ptDUP, Cdc48, Npl4, Ufd1, sPUB, sUbx, Png1, Rad23, Hsp70, PPP1), 3 Sec61 components (alpha, beta, and gamma subunit), 8 plastid division proteins (FtsZ, FtsY, MinD, MinE, MinC, ARC6/PARC6, ARC5/DRP5B, SepF), Cpn60, and PsbO. To search for the core plastid metabolism genes, core TIC/TOC components (TIC20, TIC22, TIC110, and TOC75), the SELMA components, and Cpn60, we used as queries datasets generated in (113), (16), and (15). For the remainder of the proteins, we obtained initial queries from UniProt or KEGG, and enriched these queries where necessary by using BLASTp (v.2.17.0) against our proteome database, aligning the hits alongside the queries with MAFFT v.7.526, trimming the alignments with TrimAl v.1.5.0, and constructing trees with FastTree v.2.2. These trees were then parsed to remove contaminants and paralogs, and the remaining orthologous sequences used as the enriched queries for the subsequent analyses. Subsequently, utilizing these query files, we performed BLASTp against our proteome database with an e-value cutoff of 1e-5. Collected non-*Meringosphaera* hits were passed through CD-HIT (at 85%), and then aligned together with the query files and the *Meringosphaera* hits and initial trees were constructed similarly as described above. These trees were parsed to identify deep paralogous groups, and an iterative tree-based approach using IQ-TREE version 3.0.1 was used to further remove paralogs and contaminants. Final sequences were aligned with MAFFT G-INS-i, trimmed with trimAl, and trees constructed with IQ-TREE 3.0.1 using ModelFinder. Support was calculated with a thousand optimized ultrafast bootstrap replicates. In the final trees, *Meringosphaera* sequences that branched together with at least one other *Meringosphaera* sequence from a different COSAGs were assumed to be actual *Meringosphaera* sequences and not contamination. In cases where one *Meringosphaera* sequence branched by itself, we further searched for this sequence in the individual SAGs. After identifying candidate *Meringosphaera* genes, incomplete N-termini were manually extended until the first upstream stop codon in the same frame as the start of the sequence. We then used TargetP version 2.0 (114), SignalP version 6.0 (115), and DeepLoc version 2.1 (116) to predict potential plastid targeting signal regions (i.e. transite and signal peptides). Because TargetP and SignalP only analyze the start of the sequence, we further split up our sequences into 100 bp sequences (overlapping at 50 bp) before using these prediction tools, allowing us to also investigate whether consecutive signal peptides and transit peptides were predicted.

## Data availability

Raw short-read sequence data has been submitted to the NCBI Sequence Read Archive (Bioproject: PRJNA1475611, SRR39032080-SRR39032111). 18S rDNA and 28S rDNA sequences has been submitted to GenBank (PZ506363-PZ506401 for 18S and PZ506402-PZ506453 for 28S) and the circularized *Meringosphaera* plastid genomes have also been submitted to GenBank (PZ517651-PZ517655). All code is available at https://github.com/burki-lab/Meringosphaera-SAG-analysis. Additional data is available on figshare doi:10.6084/m9.figshare.c.8543418.

## Acknowledgments

This work was supported by grants from the Swedish Research Council VR (2021-04055) to F.B and R.A.F and (2025-04150) to F.B; a European Research Council grant (ERC consolidator grant 101044505) and Science for Life Laboratory to F.B; a grant from the Gordon and Betty Moore Foundation (https://doi.org/10.37807/GBMF9201) to P.J.K; and the Knut and Alice Wallenberg Foundation Prolongation grant to R.A.F. F.B. is grateful for an International Sabbatical grant from the Faculty of Science and Technology at Uppsala University for enabling a research visit to the University of British Columbia which was determinant for this paper. We also acknowledge the Sernanders and Regnell botanical travel scholarships awarded to A.W. for helping fund the field work in Curaçao. We acknowledge the donation in Birgitta Sintring’s name to the Competence Center for Hidden Diversity in the department of Ecology and Genetics (Uppsala University) for enabling access to confocal microscopy. We thank the Hakai Institute for supporting field work on

Calvert Island and Quadra Island, and the CARBAMI foundation for supporting fieldwork in Curaçao. Sequencing was performed by the SNP&SEQ Technology Platform in Uppsala. The facility is part of the National Genomics Infrastructure (NGI) Sweden and Science for Life Laboratory. The SNP&SEQ Platform is also supported by the Swedish Research Council and the Knut and Alice Wallenberg Foundation.The data handling and analyses were enabled by resources in projects UPPMAX 2025/2-441 (and its preceding projects SNIC 2022/22-577, NAISS 2023/22-659, NAISS 2024/22-868, and UPPMAX 2025/2-160) and UPPMAX 2025/2-448 (and its preceding projects NAISS 2024/5-197 NAISS 2024/5-197, UPPMAX 2025/2-76) provided by the National Academic Infrastructure for Supercomputing in Sweden (NAISS) at UPPMAX, funded by the Swedish Research Council through grant agreement no. 2022-06725, and at Uppsala University at UPPMAX. We would like to thank the Swedish Meteorological and Hydrological Institute as well as Mikael Thollesson at the Klubban Biological Station for their help with sampling and shipment, and Elina Viinamäki and Yash Pardasani for their assistance with sample processing. We are very grateful to Megan Sørensen for designing the CARD-FISH MerChp935 plastid 16S rRNA probe.

## Author Contributions

F.B. and R.A.F. conceived this project and received the funding. P.J.K enabled field work, data collection, and contributed to discussion. A.W., F.B., and V.Z did the lab work. A.W. analyzed the data. A.W., F.B, and R.A.F drafted the manuscript. All authors edited and commented on the manuscript.

