## Supplementary material for "Single cell genomics and fluorescence microscopy suggest a permanent plastid in a marine centrohelid": SI Appendix

**Supporting Information for**
Single cell genomics and fluorescence microscopy suggest a permanent plastid in a marine centrohelid

Anne Walraven[1], Vasily Zlatogursky[1], Patrick J. Keeling[2], Rachel A. Foster[3], Fabien Burki[1]*

[1]Department of Organismal Biology, Program in Systematic Biology, Uppsala University, Uppsala, Sweden
[2]Botany Department, University of British Columbia, 6270 University Blvd. Vancouver, BC, Canada
[3]Department of Ecology, Environment, and Plant Sciences, Stockholm University, Stockholm Sweden

*Corresponding author

**This PDF file includes:**

1. Supplementary Figures S1 to S6
2. Supplementary Table S1
3. Legends for Datasets S1-S5
4. SI References

**Other supporting material for this manuscript includes:**

Datasets S1-S5

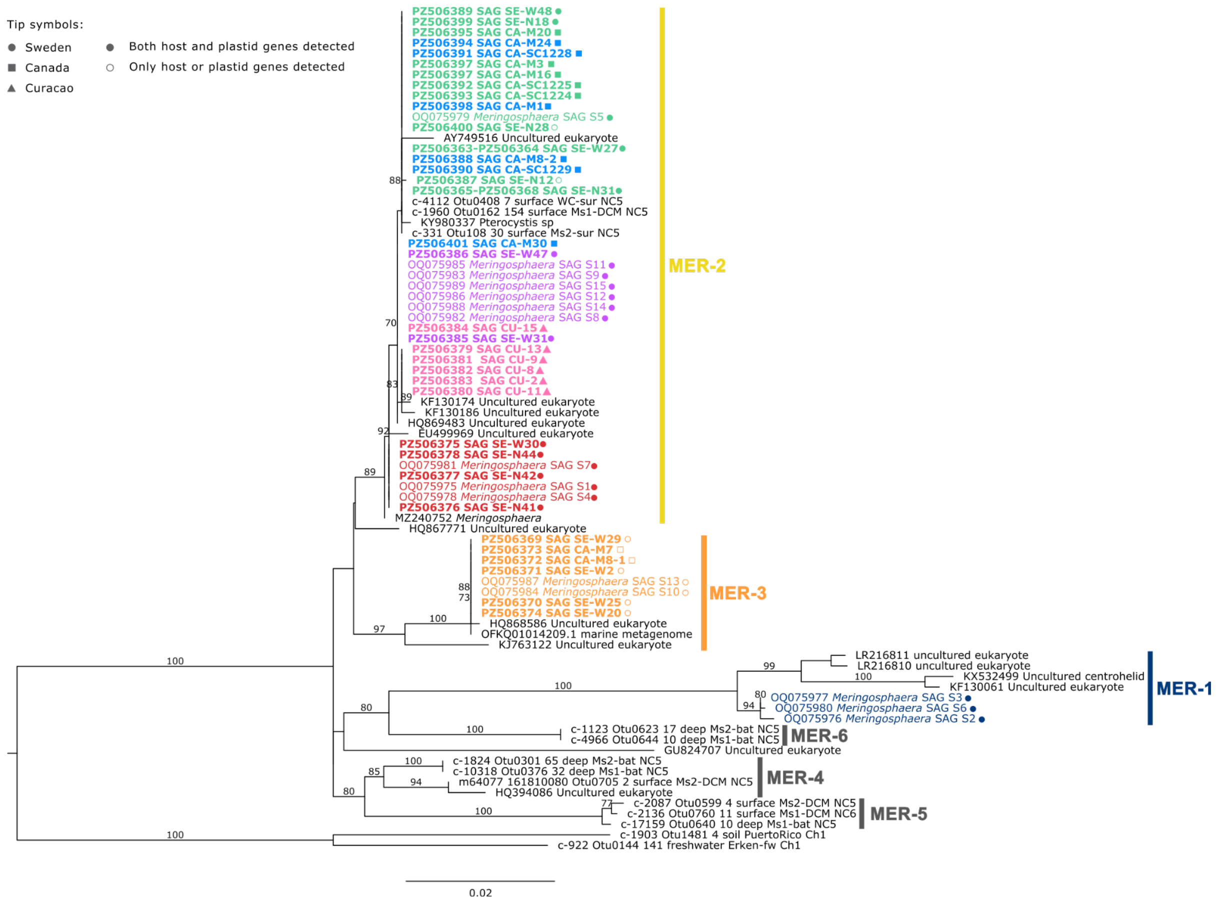


**Fig. S1.** Phylogenetic tree of *Meringosphaera* 18S rDNA, including environmental sequences. Support was calculated with a thousand ultrafast bootstrap replicates, only values equal to or greater than 70 are shown. The scale bar indicates the number of substitutions per site. Newly generated SAGs are in bold.

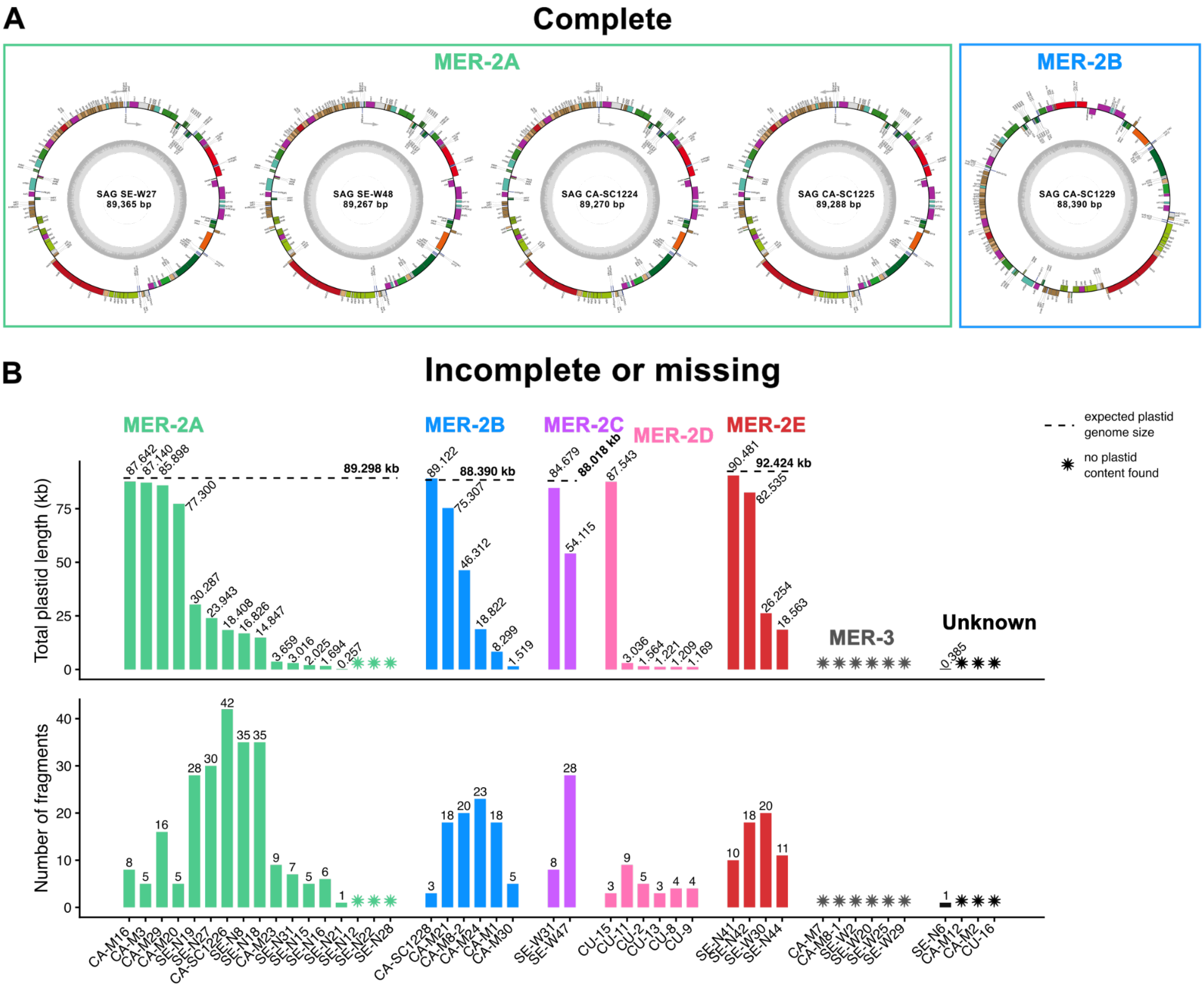


**Fig S2**. *Meringosphaera* Plastid genome assembly, including circularized plastid genomes (A) and incomplete and often fragmented or complete missing plastid genomes (B). Plastid genomes are organized by clade, or unknown if clade is not clear due to lack of data. Expected plastid genome size is based on the average size of plastid genomes in that clade, if known.

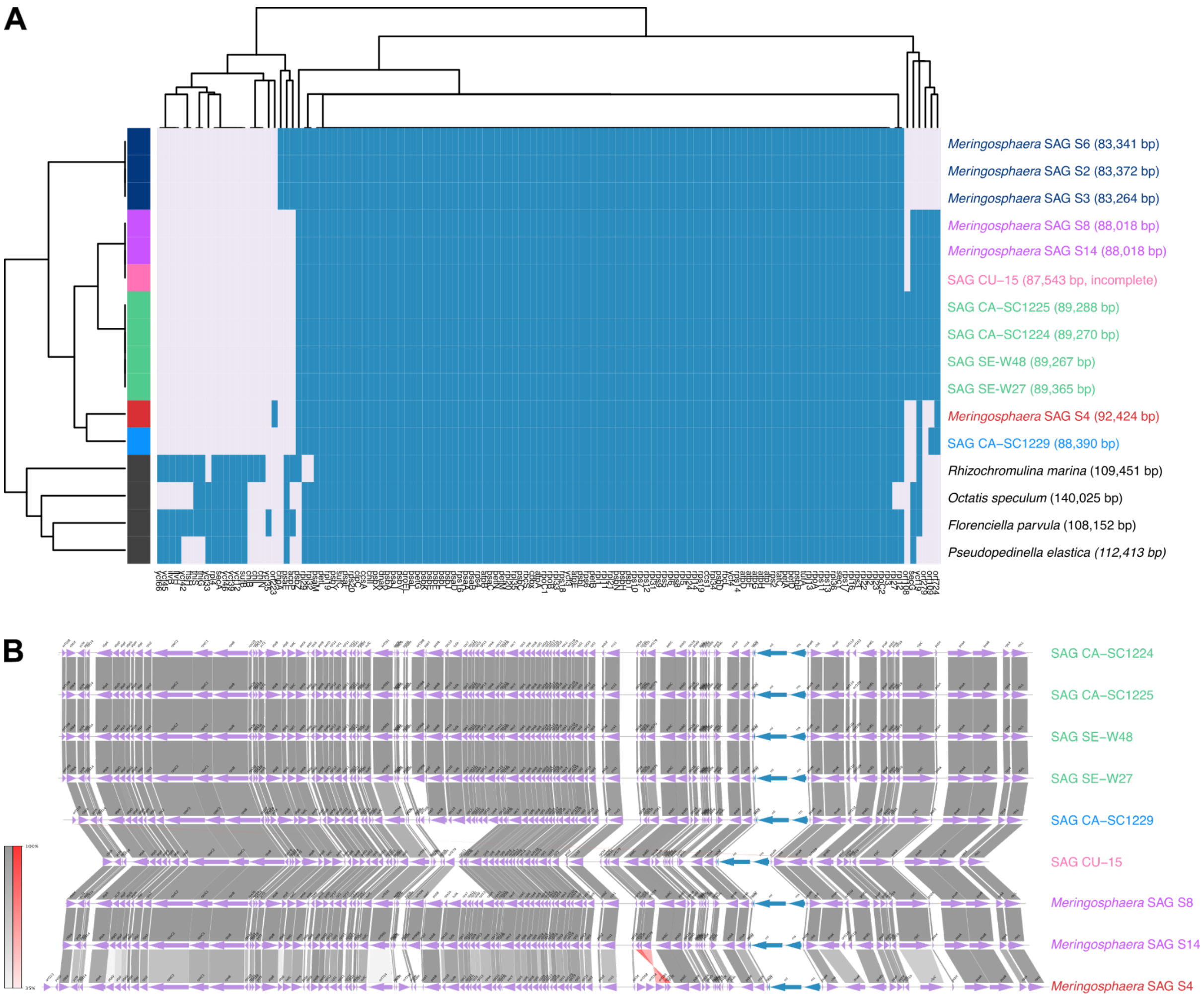


**Fig S3**. *Meringosphaera* plastid gene repertoire (A) and genome synteny (B). *Meringosphaera* plastid gene content (including *Meringosphaera* plastids previously obtained (1) is compared against gene content of other dictyochophytes (2), analyzed with Orthofinder. Present genes are shown in blue and missing genes in light purple. Plastid genome size is shown in brackets. In (B) the plastid genome synteny of MER-2 plastid genomes is shown, including all circularized plastid genomes as well as the largest fragment of the SAG CU-15 plastid genome. Plastid genomes were rotated to start at the *rbcL* gene to improve visualization. Inversions are shown in red and arrows indicate gene directionality.

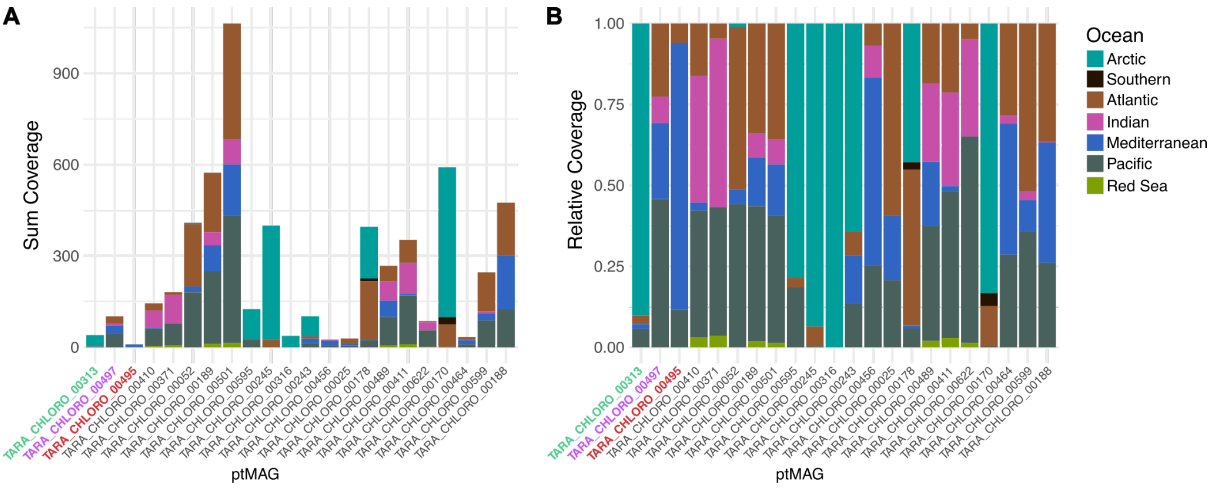


**Fig S4**. Tara Oceans plastid MAGs (3) coverage and distribution, representing all ptMAGs present in the plastid phylogenomic tree (Fig. 2). ptMAGs falling within the MER-2 clade are shown in their respective subclade color (green=MER2A, purple=MER2C, red=MER2E). Both the sum coverage (A) and relative coverage (B) are shown.

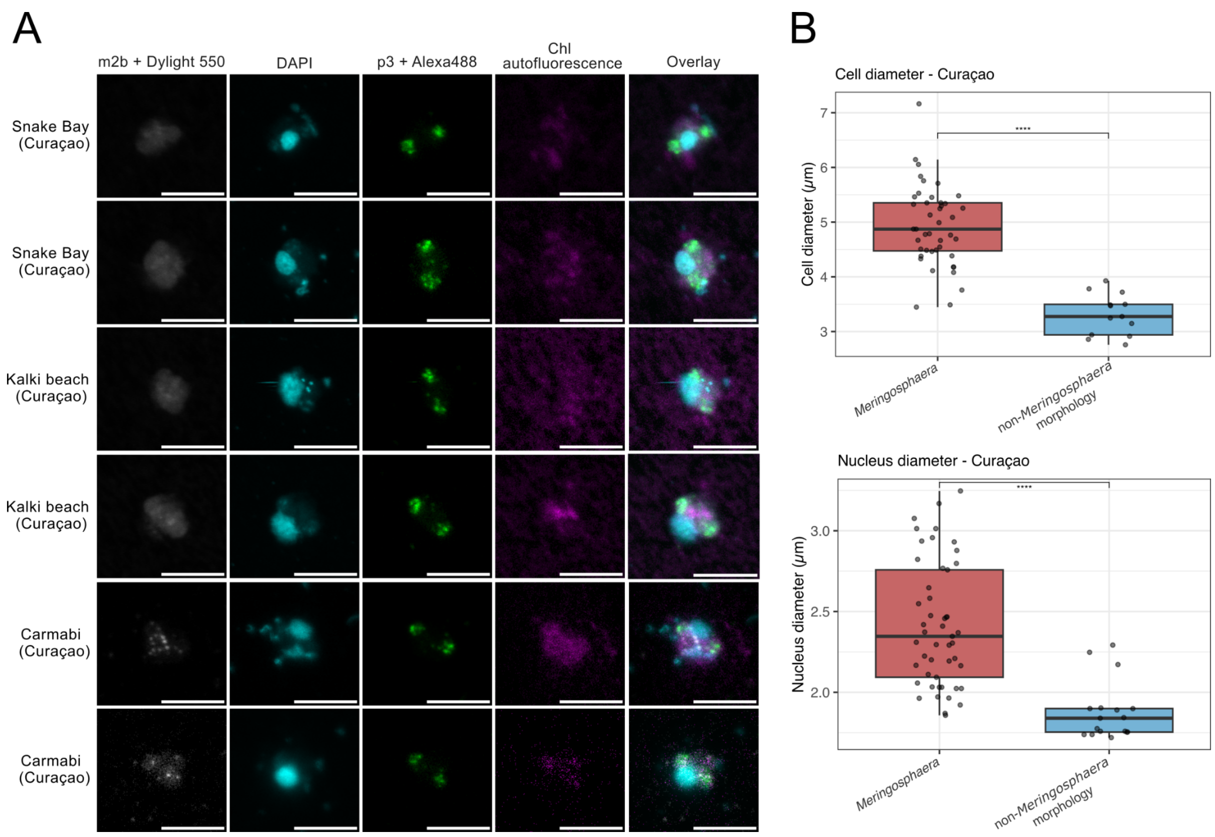


**Fig S5.** CARD-FISH of positive 16S rRNA hybridization in cells with non-*Meringosphaera* morphology (A) and comparison of *Meringosphaera* and cells with non-*Meringosphaera* morphology cell and nucleus diameter (B)**.** m2b represents the 18S rRNA probe and p3 represents the 16S rRNA probe. Scale bars represent 5 μm. For the boxplots, p-values are 2.88e-10 (cell diameter) and 2.65e-07 (nucleus diameter) calculated with the Wilcoxon rank-sum test. Only cells imaged from samples from Curaçao were compared, as that was the only location where both cell types were present.

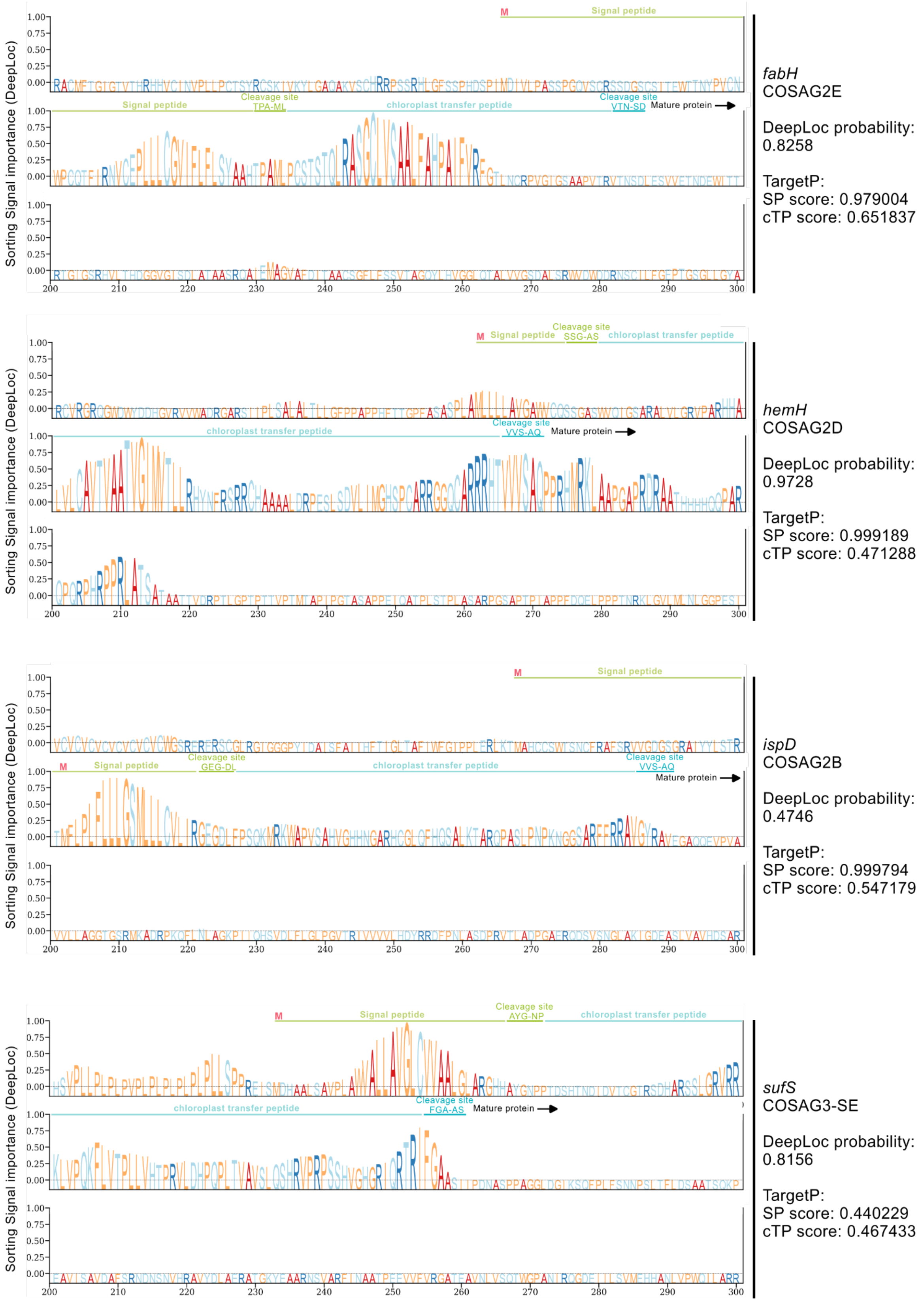


**Fig S6**. Potential bipartite plastid targeting signal in *Meringosphaera*, as predicted by DeepLoc and TargetP. Figure indicates DeepLoc plastid localization probability for the respective genes, as well as TargetP signal peptide (SP) and chloroplast transfer peptide (cTP) scores and their predicted cleavage sites. Methionines upstream of the first signal peptide cleavage site are indicated and signal peptides are extended up until that point. Only the first part of the sequence with predicted targeting peptides is shown.

| COSAG | Cells used for co-assembly | Number of contigs | Total length (Mbp) | GC-content (%) | N50 (bp) | BUSCO Complete | Fragmented | Missing |
|---|---|---|---|---|---|---|---|---|
| COSAG1 | S2, S3, S6 | 10 413 | 34 561 739 | 52.22 | 9338 | 39 | 13 | 77 |
| COSAG3-SE | S10, S13, SE-W2, SE-W20 | 30 717 | 67 779 779 | 55.78 | 3651 | 45 | 25 | 59 |
| COSAG3-CA | CA-M7, CA-M8-1 | 31 912 | 60 151 508 | 47.87 | 3111 | 11 | 12 | 106 |
| COSAG2A-SE | SE-N28, SE-W27, SE-W48 | 43 006 | 101 653 261 | 54..87 | 4733 | 47 | 21 | 61 |
| COSAG2A-CA | CA-SC1224, CA-SC1225, CA-M3, CA-M16, CA-M20, CA-M29 | 62 115 | 113 485 810 | 49.73 | 3094 | 40 | 27 | 62 |
| COSAG2B | CA-SC1228, CA-SC1228, CA-M1, CA-M24, CA-M8-2, CA-M30 | 72 497 | 128 452 016 | 51.17 | 2831 | 39 | 29 | 61 |
| COSAG2C | S8, S11, S12, S14, S15, SE-W47, SE-W31 | 30 321 | 82 013 306 | 55.78 | 6161 | 62 | 21 | 46 |
| COSAG2D | CU-2, CU-9, CU-15, CU-13, CU-8 | 183 448 | 267 067 458 | 50.26 | 1899 | 56 | 24 | 49 |
| COSAG2E | S1, S4, S7, SE-N41, SE-N42, SE-N44, SE-W30 | 39 169 | 106 095 336 | 51.90 | 5615 | 65 | 25 | 39 |

**Table S1**. COSAG statistics of the nine co-assemblies generated for this study. The table also includes which individual SAGs were co-assembled for the respective COSAG.

**Legends for Datasets S1-S5.**

**Dataset S1**. Single cell sampling metadata and SAG statististics. Metadata includes including isolation location, coordinates, sampling date and isolation date, and the presence of chlorophyll autofluorescence when an epifluorescent microscope was available. If known, respective clades each cell belongs to are shown, which were inferred for either host or plastid phylogenies, or both. SAG statistics includes genome statistics of the 50 SAGs generated for this project, after cleaning with Blobtools, as well as 18S and 28S rDNA sequence length and accession. Where multiple accessions are shown, fragments were merged for phylogenetic analysis.

**Dataset S2**. Sampling metadata for the samples used for CARD-FISH, as well as 16S rRNA probe scoring, chlorophyll scoring, and cell and nucleus diameter for all CARD-FISH cells imaged. Metadata includes not only location and sampling date, but also fixation method (where samples first filtered and and then fixed, or first fixed and then filtered), filter pore size, and sampling volume.

**Dataset S3**. File containing all taxa used alongside the nine Meringosphaera co-assemblies in the search for host-encoded plastid-associated genes.

**Dataset S4**. File containing all Meringosphaera candidate host-encoded genes involved in plastid metabolism and Cpn60 found in this study, alongside their top Swissprot hit and predicted localization and predicted targeting/signal peptides from DeepLoc, TargetP, and SignalP.

**Dataset S5**. Method specification for all the SAGs generated for this study as well as those published previously (1), where there were differences in their treatment.
